# Effective concentrations enforced by intrinsically disordered linkers are governed by polymer physics

**DOI:** 10.1101/577536

**Authors:** Charlotte S. Sørensen, Magnus Kjaergaard

**Affiliations:** Department of Molecular Biology and Genetics, Aarhus University; The Danish Research Institute for Translational Neuroscience (DANDRITE); Aarhus Institute of Advanced Studies (AIAS); The Center for Proteins in Memory (PROMEMO)

## Abstract

Many multidomain proteins contain disordered linkers that regulate inter-domain contacts, and thus the effective concentrations that govern intra-molecular reactions. Effective concentrations are rarely measured experimentally and therefore little is known about how they relate to linker architecture. We have directly measured the effective concentrations enforced by disordered protein linkers using a new fluorescent biosensor. We show that effective concentrations follow simple geometric models based on polymer physics, offering an indirect method to probe the structural properties of the linker. The compaction of the disordered linker depends not only on net charge, but also on the type of charged residues. In contrast to theoretical predictions, we found that polyampholyte linkers can contract to similar dimensions as globular proteins. Hydrophobicity has little effect in itself, but aromatic residues lead to strong compaction likely through π-interactions. Finally, we find that the individual contributors to chain compaction are not additive. This work represents perhaps the most systematic study of the relationship between sequence and structure of intrinsically disordered proteins so far. A quantitative understanding of the relationship between effective concentration and linker sequence will be crucial for understanding disorder-based allosteric regulation in multidomain proteins.

## Introduction

Protein interactions are tightly regulated. The specificity is not only determined by the protein structure, but also by which molecules a protein encounters. Molecular encounters are often enhanced by co-localization or even a direct physical link between the interaction partners. Such links are created by hundreds of anchoring and scaffolding proteins that connect other molecules.^1–3^ Furthermore, multidomain proteins often contain long linkers that play a similar role for intra-molecular interactions.^4^ A physical connection often increases encounter rates by several orders of magnitude, which shifts equilibrium position of binding reactions and the rates of biochemical reactions by a similar amount.^5,6^ The encounter frequency of such linked reactions is concentration independent. Instead it depends on the architecture of the connection, and therefore linker properties directly affect protein function.

Anchoring proteins and interdomain linkers often belong to the family of intrinsically disordered proteins (IDPs).^7^ Disordered linkers allow the tethered domains to contact in any orientation, and therefore they represent a generic mechanism for fusing domains, which cannot be accomplished by rigid proteins. The architecture of the connection between molecules is not static, but can vary in time and space to regulate the functions of the tethered domains. Recently, it has been proposed that the concepts of allostery needs to be updated to include structural disorder: Here allostery is transmitted through changes in heterogenous ensembles rather than through structured domains.^8,9^ Most structural changes occurring in linker regions will affect the function of the tethered domains. Therefore, linkers provide a generic mechanism whereby an event occurring in one part of a protein can affect distant parts. This is the defining feature of allostery. To understand such allosteric effects, it is crucial to study the role of linkers quantitatively.

The functions of linkers can be understood quantitatively in terms of effective concentrations. For an intra-molecular reaction, the encounter rate between tethered domains equals the rate of the same untethered reaction at a given concentration.^10^ This concentration is known as the effective concentration. Formally, the effective concentration is defined as the ratio of the equilibrium constants for two equivalent binding reactions, where one occurs intra-and one inter-molecularly.^11^ When the linker is sufficiently long to join the binding sites without strain, the effective concentration is independent of what is linked and solely a property of the linker.^12,13^ Intriguingly, this suggests that effective concentrations can be measured in a convenient model system and extrapolated to other systems. Effective concentrations can be measured by competition experiments, where a free ligand displaces a tethered ligand.^11^ Such measurements have mostly been used in efforts to optimize multivalent drugs through avidity.^14,15^ In molecular biology, effective concentrations are rarely measured experimentally, but usually estimated theoretically from the volume in which the tethered ligand is free to diffuse.^16–20^ While such simple geometric models are commonly used, they have not been tested in complex biological systems.

The simplest linker is a fully disordered chain. Fully disordered linkers are an attractive model for understanding effective concentrations as they are well-described by theories borrowed from polymer physics. When the sizes of disordered proteins are measured as e.g. end-to-end distances or the radius of hydration or gyration, it scales with the chain length following a power law such as:

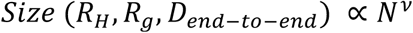

Where N is the number of residues and *v* is a scaling exponent determined by chain compaction. Such scaling laws underpin theoretical calculations of effective concentrations and depends on estimating the scaling exponent *v*. On average, IDPs have been found to have *v* values from 0.51-0.58,^21–23^ but the scaling exponents of disordered proteins varies from about 0.4 for disordered states of foldable proteins to about 0.72 for highly charged IDPs.^24^ For reference, globular proteins and rigid rods have scaling exponents of 0.33 and 1, respectively. The sequence-compaction relationship of IDPs has been studied by correlating chain size with variations in sequence. Net charge dominates chain compaction through intra-chain repulsion^21,24–26^. Furthermore, compaction is weakly correlated to hydrophobicity and weakly anti-correlated to proline content.^21^ There is a discrepancy in the literature regarding the effect polyampholyte strength as it has been predicted to cause both compaction or expansion.^27,28^ Furthermore, the patterning of charged residues is also critical,^28–30^ which vastly increases the potential complexity.

Here we investigate how effective concentrations in multi-domain proteins depend on linker architecture. We directly measure effective concentrations for many disordered linkers with systematic changes in the physical properties of the linker. Our new fluorescent biosensor for measurement of the effective concentrations provides a new way to probe sequence-compaction relationships in intrinsically disordered proteins and relating these to biochemical function.

## Materials and methods

### Preparation of DNA constructs

DNA constructs were obtained from Genscript by insertion of synthetic genes between the NdeI and BamHI sites of a pET15b vector, and sub-cloning of new linkers using unique NheI and KpnI sites flanking the linkers. Full protein sequences are given in the supplementary materials.

### Protein expression and purification

All fusion protein constructs were expressed in BL21(DE3) cells in 50mL ZYM-5052 auto-induction medium^31^ supplied with 100μg/mL ampicillin and shaking at 120 RPM. The temperature was kept at 37°C for 3 h, and thereafter decreased to 18°C. The cells were harvested by centrifugation (15min, 6.000g) after 40-48 h, when the cultures had changed color indicating mature fluorescent proteins. Bacterial pellets were lysed using B-PER Bacterial Protein Extraction Kit (Thermo Scientific) according to manufacturer’s protocol, and the lysate was applied to gravity flow columns packed with nickel sepharose. After washing with 20 mM NaH_2_PO_4_ pH 7.4, 0.5 M NaCl, 20 mM imidazole, fusion proteins were eluted by increasing the imidazole concentration to 0.5 M. Fusion proteins were subsequently purified using Strep-Tactin XT Superflow (IBA) columns according to the manufacturer’s instructions, and dialyzed overnight into Tris buffered saline (TBS). The MBD2 peptide was expressed overnight in BL21(DE3) cells at 37°C in ZYM-5052 auto-induction medium with 100μg/mL ampicillin and shaking at 120 RPM. The cells were resuspended in 20 mM NaH_2_PO_4_ pH 7.4, 0.5 M NaCl, 20 mM imidazole and lysed by heating to 80 °C for 20min,^32^ and debris pelleted by centrifugation (15min, 14.000g). The peptide was purified by nickel sepharose as above except a stepwise elution up to 0.5 M imidazole was used before dialysis into TBS. It was critical to prepare and concentrate the MBD2 peptide freshly and store it on ice until use. Protein concentrations were measured using A_280_.

### Measurement of effective concentrations

0.1 μM of each fusion protein was titrated with the MBD2 peptide through 16 serial two-fold dilutions in TBS. The starting concentration was in the range of 1.6-2 mM for WT MBD2 and 3.3 mM for the V227A MBD2 mutant. Samples were analyzed in triplicate in black 386-well plates with 1g/L bovine serum albumin (BSA) (Fisher Scientific) added to prevent sticking. The FRET measurements were performed in a SpectraMax I3 platereader using donor excitation at 500nm, and emission detected in 25nm-wide bands centered at 535 and 600 nm. The titration data was analyzed by non-linear fitting in MATLAB to the standard fitting equation for 1:1 binding reactions with K_d_ replaced by c_e,app_:

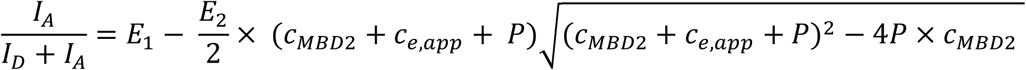

For titration with the WT MBD2 peptide, this determines an “apparent effective concentration”, which was multiplied by the affinity ratio of the wild-type and mutant peptides to produce the true effective concentration. The correction factor was established to be 30 by titration of the fusion protein containing the GS_120_ linker with the V227A MBD2 peptide.

## Results

### Reporter design

We designed a fusion protein inspired by FRET biosensors^33^ to measure effective concentrations for different linker architectures. An exchangeable linker joins two protein domains that form an interaction pair. These domains are flanked by the fluorescent proteins mClover3 and mRuby3, which form a FRET pair^34^ (Fig. 1A). When the intra-molecular complex is formed, the fluorescent proteins are brought into close contact resulting in efficient FRET. The effective concentration is measured by following the FRET efficiency in a titration,^11^ where a free ligand displaces the intra-molecular interaction. The ideal interaction pair is small, has a known 3D structure, is easy to produce in *E. coli* and binds tightly to ensure full ring-closing. Furthermore, in the bound state the N-terminus of one protein should be close to the C-terminus of the other and *vice versa*. This ensures a high FRET efficiency in the closed state and allows even a short linker to join the domains without strain. These constraints are ideally fulfilled by an anti-parallel heterodimeric coiled-coil such as the complex between MBD2 and p66α used here (Fig. 1B).^35^

**Figure 1:**
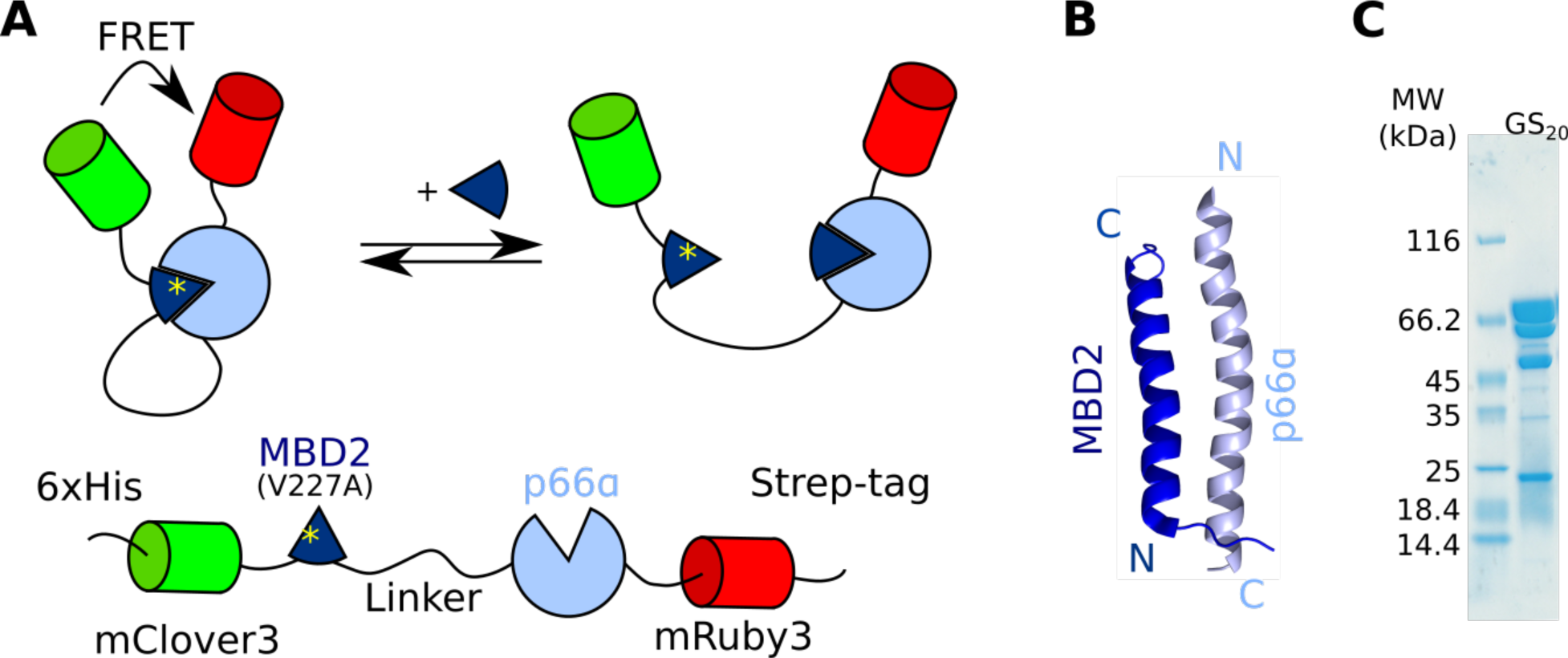
A FRET biosensor for measuring effective concentrations. A) Titration with free MBD2 peptide displaces the intra-molecular interaction and results in a decrease in the FRET efficiency. B) Structure of the interaction pair in the fusion protein : The anti-parallel coiled-coil formed between MBD2 and p66α (PDB:2L2L)^35^ C) SDS-PAGE gel of the purified fusion protein reveals internal cleavage sites in mRuby3. The cleavage products cannot be removed, but do not affect measurements.

The fusion proteins have purification tags at both the N-and C-termini to allow parallel purification of many constructs by sequential affinity chromatography. After purification, SDS-PAGE revealed a major band corresponding to the expected molecular weight of the fusion protein (Fig. 1C). The four minor bands corresponded to proteolytic cleavage in the mRuby3 domain as revealed by mass spectrometry. These bands could not be removed by gel filtration. This suggests that the cleaved fluorescent proteins remain in a stable complex, although it is not clear whether it is fluorescent. A fraction of inactive fluorophores will decrease the FRET amplitude, but will not affect the mid-point of the titration and the measurements of effective concentrations.

### Measurement of effective concentration

Fusion proteins were initially constructed with linkers consisting of (GS)_n_ repeats ranging from 20 to 120 residues. A 240-residue GS-linker was also tested, but resulted in insoluble protein. Titration of all constructs with free WT MBD2 peptide resulted in a sigmoidal decrease of the proximity ratio (Fig. 2A) where the titration midpoint moved to lower concentrations with increasing linker length. Simultaneously, the proximity ratio of the post-titration baseline decreased with linker length consistent with a more expanded open form (Fig 2A). Across the whole dataset, the pre-and post-transition proximity ratios varied unsystematically, which we believe was due to small differences in fluorophore maturation. This complicated the determination of intra-molecular distances from FRET values, and therefore we only extracted the midpoint of the titration.

**Figure 2:**
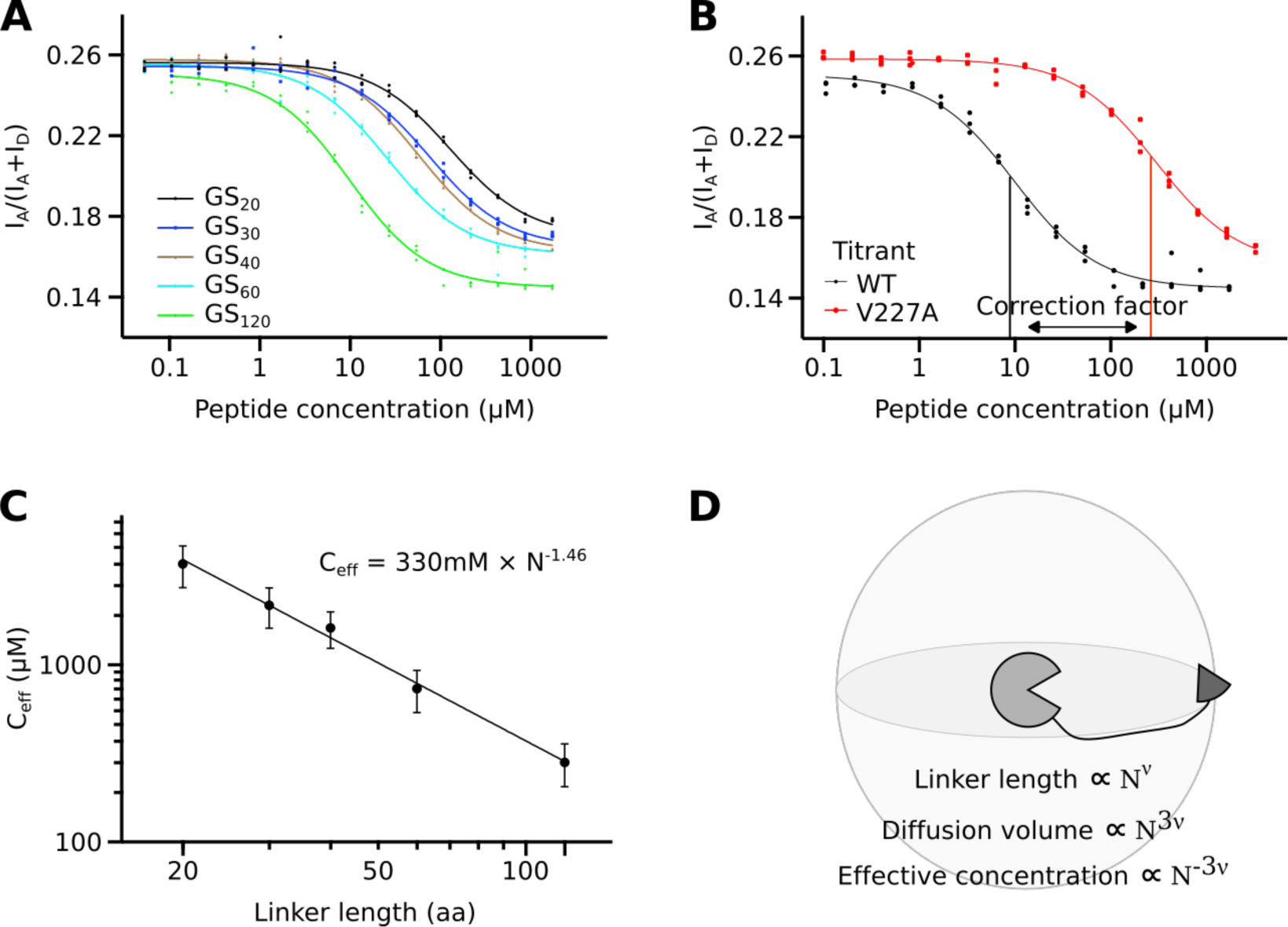
Measurement of effective concentrations. A) Titration curves of fusion proteins containing GS-linkers of variable length reveal that the effective concentration scale monotonously with linker length. B) The correction factor corresponding to the difference in affinity between WT and V227A was determined by titration of GS_120_ with both peptides. C) Effective concentrations follow a power law as revealed by the straight line in a log-log plot. The scaling exponent which is the focus of the remainder of the manuscript is derived from non-linear fitting as indicated. D) The expected geometric relation between scaling exponents for protein size, *v*, and effective concentration. The linker defines a volume in which tethered ligand can diffuse, and the effective concentration is inversely proportional to this diffusion volume.

Fitting of titration data requires the concentration of the free ligand to exceed the mid-point by a factor of ten. This is impractical for effective concentrations that were expected to reach the millimolar-range. Therefore, the fusion proteins contained a weakened mutation of MBD2, and were titrated with WT MBD2 peptide. This shifted the midpoint by the ratio between WT and mutant affinities, which subsequently was applied as a correction factor. To determine the correction factor, we titrated the fusion protein with the GS_120_ linker with both WT and mutant peptide (Fig. 2B). The titration midpoint was shifted by a factor of 30, which was applied to the titration midpoint of all variants to produce the effective concentration. As it was impractical to prepare competitor peptides for each linker composition, we used the titrant peptide with a flanking GS-segment for all other linker compositions. As the flanking linker residues may affect the stability of the complex, the correction factor may differ for other linker compositions. A mis-matched correction factor results in a constant shift of the polymer scaling law, but should not affect the scaling exponent.

The effective concentration scales with linker length following a power law as shown by the straight line in Fig. 2C. Notably, this conclusion does not require any assumptions of the distribution of the linker conformations. Relative to one binding partner, geometric considerations suggest that the volume accessible to the other partner scales with the linker length with an exponent of 3*v* (Fig. 2D). Accordingly, the effective concentrations should scale with an exponent of −3*v*. Fitting of effective concentrations from the GS-linker series gave a scaling exponent of −1.46 corresponding to a *v* of 0.49. The GS-linker is a polar tract in a recent systematic classifications,^28^ and is thus expected to form a relatively compact globules due to backbone interactions.^36^ Accordingly, the GS-linker was slightly more compact than denatured proteins^37^ and IDPs.^21–23^ The excellent agreement with a power law and scaling exponent of approximately −3*v* suggested that it is valid to estimate effective concentrations based on geometric considerations.

### Variation of the linker sequence

The excellent agreement with a power law suggested that effective concentrations could be used to probe the sequence-compaction relationship of IDPs. To probe how the effective concentration depends on linker sequence, we systematically varied the properties that were likely to affect linker compaction: charge, ampholyte strength, rigidity, hydrophobicity and aromaticity. We systematically increased the net charge of the linker by introduction of charged residues into a GS-linker. All linkers used here have a uniform pattern through-out the sequence (Fig. 3A), which allows linkers of different lengths to be described by a single scaling exponent. For each linker composition, we generated linkers with a total of 20, 40, 60 and 120 residues, and measured the effective concentrations through titration experiments. In a few case near the limit of solubility, the longest linker was shortened to 80 residues or the series only contained the three shorter linkers (Table S1). Each linker composition followed a power law as illustrated for linkers containing glutamate residues (Fig. 3B, Fig. S1). Both the pre-factor and the scaling exponent from the fit varied with linker composition. The variation of the pre-factor may be due to mis-match of the correction factor, so in the following we concentrate on the scaling exponent that reports on linker compaction.

**Figure 3:**
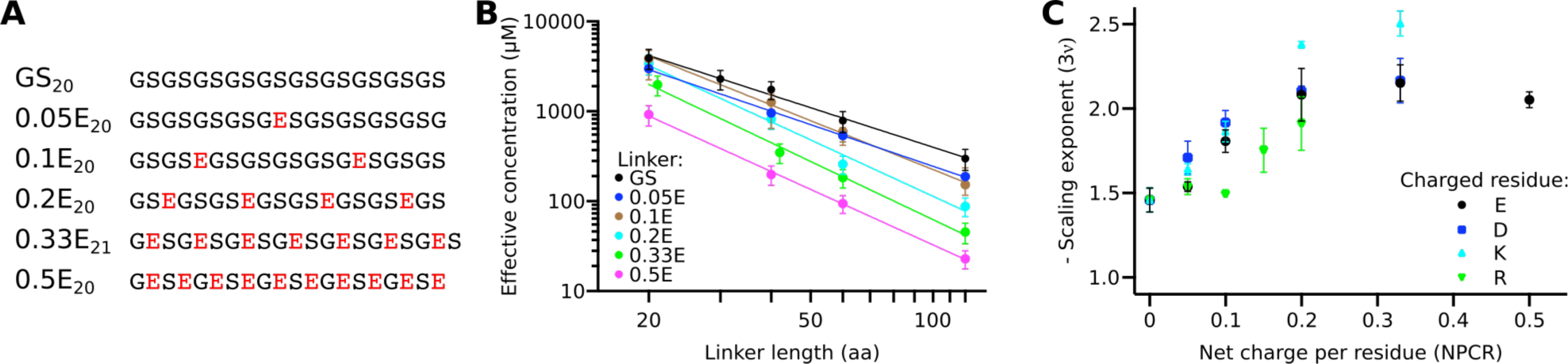
Net charge per residues dominates the scaling of effective concentration with linker length. A) Net charge was uniformly distributed into a background of GS-linkers. The same pattern is used for subsequent linker-series B) Measurement of effective concentration allows determination of scaling exponents for each linker composition. C) Scaling exponents demonstrate a linker expansion with increasing net charge. Compared to the negatively charged residues, arginine expands the chain less and lysine more.

### Are all charged residues equal

To test if all charged residues affect linker compaction equally, we measured effective concentrations for linkers containing increasing amounts of the four charged residues. For each residue-type, increased net charge per residue lead to chain expansion (Fig. 3C). Glutamate and aspartate caused an equal expansion: The scaling exponent changed gradually from −1.46 (*v*=0.49) in the uncharged linker to ∼-2.1 (*v*=∼0.7) at a net charge per residue of 0.2. The scaling exponent did not increase further with increasing net charge suggesting an upper limit that linker does not expand beyond. This limit corresponds to the *v* previous observed for highly charged IDPs.^24^ Lysine-containing linkers initially followed the expansion of negatively charged linkers, but continues up to a scaling exponent of ∼-2.4 (*v*=∼0.8). This is higher than the *v*-values reported for any other disordered protein. In contrast, arginine-containing linkers have the opposite effect: Residue fractions up to 0.1 had the same scaling exponent as GS-linkers. At higher fractions, the scaling exponent increased, but did not reach the same expansion as the other residue types even at the highest fractions permitted by solubility. This demonstrated that charged residues types have a different impact on IDP compaction.

### Polyampholyte linkers

Most IDPs are polyampholytes, which entails that chain compaction is determined by the balance between attractive and repulsive interactions. To investigate the effect of ampholyte strength, we created linkers with equal numbers of glutamate and arginine residues. At low fractions of charged residues, the scaling exponent was roughly constant, but there appeared to be a threshold after which the chain contracts dramatically (Fig. 4A). The last data point at a fraction of charged residues of 0.67 suggested a scaling exponent of −1 (*v*=0.33), which is the same compaction as globular protein. This observation agrees qualitatively with the contraction caused by screening of charges in a polyampholyte,^38^ but contradicts the diagram-of-states representation of IDP classes.^28,29^

**Figure 4:**
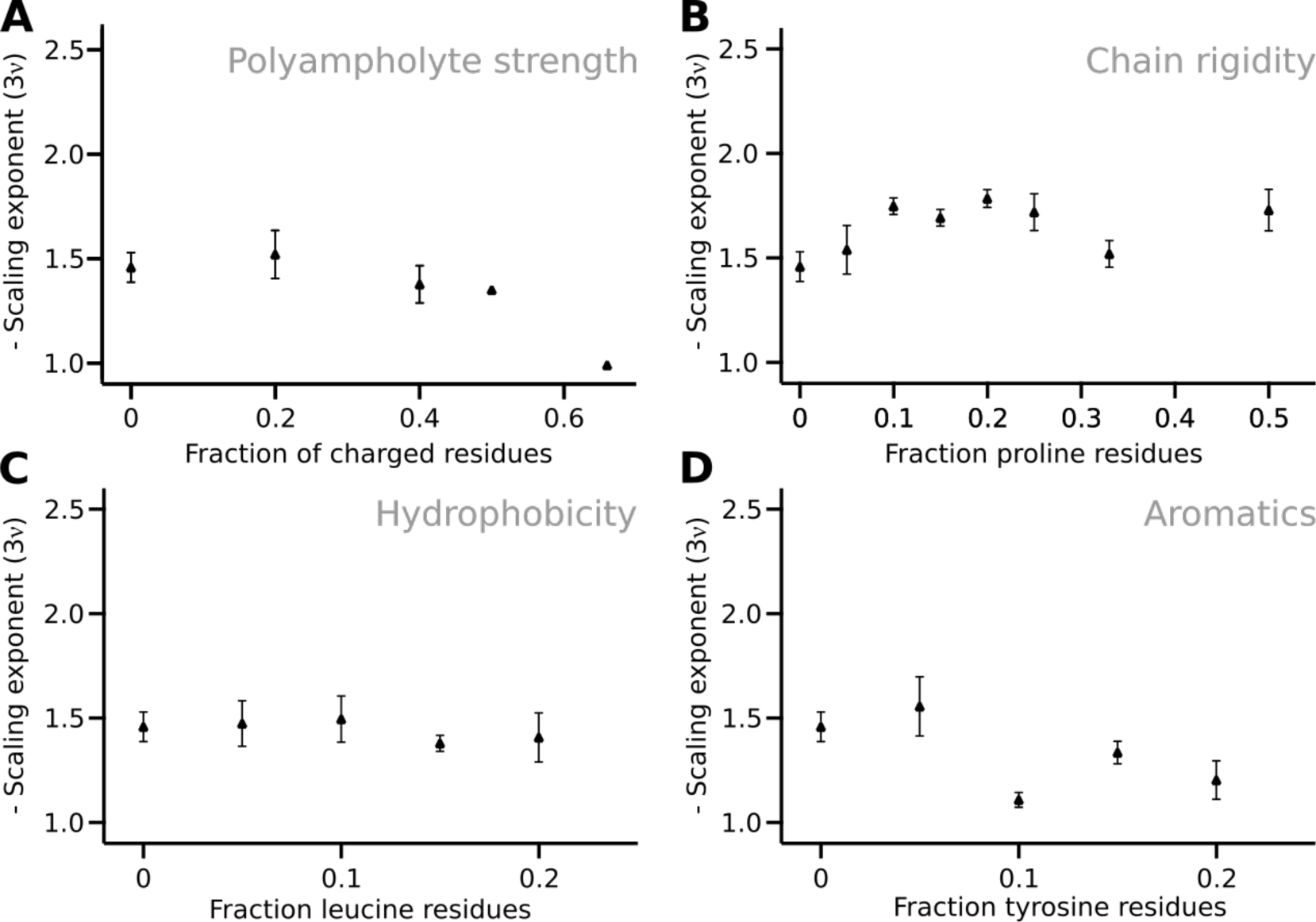
The effect of polyampholyte strength, proline, leucine and tyrosine residues on scaling exponents. A) Scaling exponents from neutral polyampholyte linkers with an equal proportion of arginine and glutamate residues show a strong compaction to a globular chain. B) Chain expansion shows a complex dependence on proline content. At low fractions of proline residues, proline residues lead to linker expansion, which may be reverse at high proline fractions. C) Hydrophobicity was increased by introduction of leucine residues, but lead to practically no change in chain compaction. D) Tyrosine residues lead to strong compaction, likely due to *π*-interactions. Error bars are 95% confidence intervals estimated from the fit.

### The effect of chain rigidity

The rigidity of IDP backbone is mainly determined by the fraction of proline and glycine residues. The baseline GS-linker is highly flexible, and we therefore increased rigidity by introducing proline residues (Fig. 4B). The scaling exponent increased to a plateau at ∼-1.7 (*v* = ∼0.57), which is reached at a proline fraction of 0.1, which is similar to the dimensions of chemically denatured proteins.^37^ Proline thus expanded the linker to a lesser extent than charged residues in agreement with previous studies.^21^ Unlike the other series tested here, the expansion with increasing proline content appeared non-monotonic. This could suggest that different effect dominate at different fractional content of proline, e.g. the propensity to form polyproline II helices.

### The effect of hydrophobic interactions

It is unclear strongly hydrophobicity affects compaction of disordered proteins. In IDPs, hydrophobicity is weakly anti-correlated with *v*,^21^ whereas in disordered states of foldable proteins this anti-correlation is strong.^24^ To assess the effect of hydrophobicity systematically, leucine residues were introduced into the linker. We chose leucine as it is among the most hydrophobic of the non-aromatic residues,^39^ but is not *Β*-branched and thus less likely to perturb backbone dihedral distributions. Due to solubility, the linker sequence can only be extended up to leucine fractions of 0.2. Introduction of leucine residues lead to a small decrease in scaling exponent to −1.4 (*v*=0.47) (Fig. 4C). This qualitatively agreed with a weak anti-correlation between size and hydrophobicity.

### π-interactions between aromatic residues

Interactions between *π*-electrons in aromatic side chains have a strong potential to induce intra-chain interaction, most recently demonstrated by their effect on liquid-liquid phase-separation.^40^ We introduced tyrosine residues as it is the least hydrophobic aromatic amino acid and has the largest effect on phase separation.^40^ Tyrosine residues caused a noticeable reduction in the scaling exponents already at a fractional content of 0.1, where the scaling exponent was ∼-1.2 (*v*=0.4) (Fig. 4D). The contraction was smaller than the contraction observed for polyampholytes, although the fractional content of aromatic residues cannot be increased as far due to insolubility. However, the effect of tyrosine residues was larger than that of leucine, which suggests that *π*-interactions are more important to chain compaction than hydrophobicity.

### Are contributions to chain compaction additive

Previous studies have identified individual factors that affect IDP compaction, but have not clarified whether they are additive. We therefore constructed linkers simultaneously increasing net charge and proline content. When probed alone, proline expanded linkers measurably already at fraction of 0.05. However, when proline was introduced together with a charged residue, it did not lead to further expansion, simply resulted in identical scaling exponents as glutamate-only linkers (Fig. 5A). This suggested that expansion caused by chain rigidity and net charge are not additive. Instead, the mixed sequences were simply dominated by the strongest individual effect.

**Figure 5:**
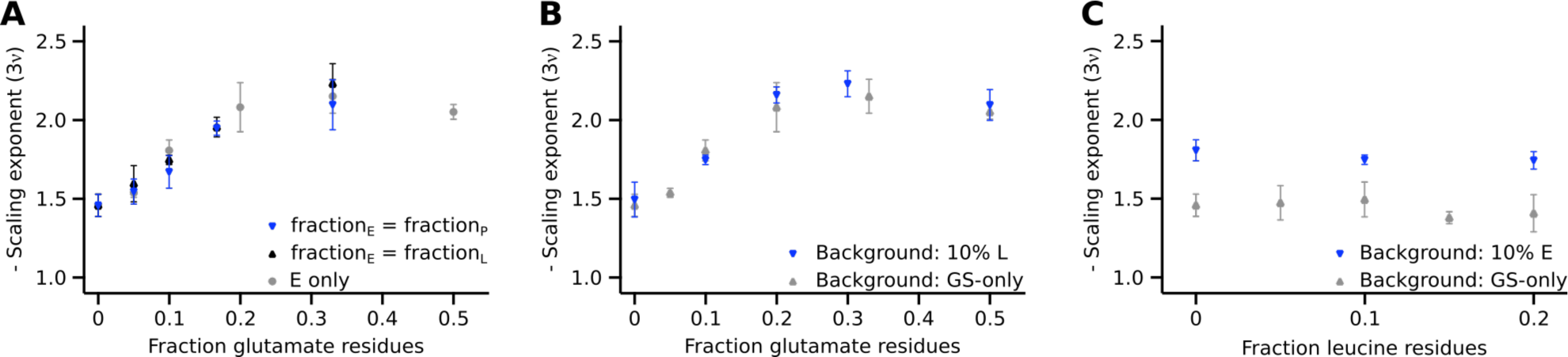
Lack of additivity of individual contributions. A) Proline or leucine residues were introduced in equal proportion to glutamate residues, but the scaling coefficient follows that of the glutamate only series. This suggests that the expansion caused by net charge and chain stifness are not additive. b) Charge expansion caused by glutamate against a constant fraction of 0.1 of leucine residues shows that hydrophobicity has a negligible effect. c) Increasing fractions of leucine only has a negligible effect on the dimensions of the linkers expanded by charge.

### Does hydrophobicity modify charge expansion

Hydrophobicity had a surprisingly small effect when investigated alone. To further test this conclusion in the context of additivity, we created mixed linker series containing glutamate and leucine residues in equal proportion. Again, the scaling exponent followed that of charge series alone. This series changed two parameters at once, so we designed new linker series where either the glutamate content or the leucine content was varied and the other held constant at a residue fraction of 0.1. When the charge expansion was probed in the context of a fraction of 0.1 leucine residues, the curve again followed the glutamate-only series (Fig. 5B). When the fraction of leucine residues was increased within the limits permitted by solubility, the scaling exponent decreases slightly, although within error (Fig. 5C). In total, these data suggest that hydrophobicity in itself has a vanishing effect on the compaction of IDPs, at least within the limits of protein solubility. Furthermore, the factors that affect IDP compaction are not necessarily additive.

## Discussion

Linkers control many biochemical reactions via the effective concentration. Here we have shown that effective concentrations in multidomain proteins with disordered linkers follow polymer scaling laws. We have thus experimentally validated the geometric models commonly used to estimate effective concentrations, but also show that the effective concentration depends strongly on linker sequence. For a 100-residue linker, the difference between *v* values of 0.4 and 0.7 corresponds to a 63-fold change in effective concentration. This can be the difference between an intra-molecular interaction being saturated or hardly formed at all. Changes in the linkers following e.g. ligand-binding or post-translational modification may thus be directly transmitted into allosteric regulation of the domains tethered at the end. Linkers may thus be one of the most direct examples of how the structural properties of intrinsically disordered proteins affect biochemical function, underscoring the need to understand the relationship between IDP sequence and compaction. In the short-term, direct measurement of the effective concentrations using the system developed here may help us understand allostery in IDPs. ^9^

### Sequence-structure relationships in IDPs

Polymer models have been used successfully to describe the structural properties of IDPs and are the foundation for theoretical predictions of effective concentrations. Here we show that the relationship can be reversed: Measurement of effective concentrations is an efficient way to parametrizing polymer descriptions of IDPs and describe the relationship between sequence and compaction. While we study interdomain linkers, it is likely that the conclusions can be generalized to other types of IDPs.

### Net charge

Net charge is the strongest predictor of the compaction of IDPs.^21,24,25^ Previous studies suggest that the maximal expansion is reached at a net charge per residue of ∼0.4. In contrast, we find a value of ∼0.2 (Fig. 3C). In previous studies, the charge expansion occurred against a complex background sequence, and there may thus be compensating attractive interactions. Our value may thus represent a pure estimate of how much charge density it takes to fully expand an otherwise inert IDP, whereas the value of around 0.4 may be more relevant for complex sequences with additional attractive interactions. Another key difference is the role of arginine residues. Arginine form attractive interactions that partially the charge-charge repulsion. This recapitulates the role of arginine in self-association of IDPs during liquid-liquid phase separation, and may thus be due to its capacity to form *π*-interactions.^40^ Conveniently, for most positively charged protein sequences this effect may be offset by the higher than average repulsion of lysine residues. Therefore, chain properties may be well-described in terms of the net charge density as long as lysine and arginine are equally common.

### Polyampholyte sequences

The compaction of polyampholytes is determined by the balance between attractive and repulsive interactions. As controlled experiments on polyampholytic IDPs have been scarce, organic polyampholytes have been used as models to understand the impact of ampholyte strength on IDP structure. Organic polyampholytes form a range of compact structures, and at high charge densities they eventually become almost globular albeit with a liquid-like internal structure.^41^ It is not clear how well such models describe proteins. In one case, increase of the ionic strengths lead to expansion of a polyampholytic protein,^38^ which suggested that polyampholyte interactions are overall attractive. This conclusion is also supported by the compact state adopted by the complex between two oppositely charged IDPs.^42^ On the other hand, a computational study of polyampholyte sequences suggested that the compact state only arises if charges are unevenly distributed, whereas a well-mixed polyampholytes was predicted to form expanded coils.^29^ This is summarized in the diagram-of-states description of IDPs, where increase of the strength of neutral polyampholyte leads to a globule-to-coil transition.^28^ In contrast, we find that increase of the polyampholyte strength lead to compaction (Fig. 4A), which eventually reaches the same dimensions as a globular protein. One possible explanation for this discrepancy is the difference between arginine and lysine. The computational study used lysine,^29^ whereas we used arginine. Whether ampholyte interactions are attractive or repulsive may thus depend on the type of positive amino acids.

### Hydrophobic side chains in IDPs

Hydrophobic side chains in IDPs are mostly solvent exposed. Contraction may bring such side chains in proximity to interact and thus form a partial protection from the solvent. Therefore, hydrophobicity has been found to be anti-correlated with *v* in disordered proteins.^21,24^ By testing this finding in a variety of sequence contexts, we found that increasing the hydrophobicity did not lead to a noticeable contraction of the linker. One explanation for this is that the disorder of the chain prevents the proteins from forming even partially desolvated hydrophobic interactions. In complex sequences, hydrophobicity correlates with other factors that could cause chain compaction. Such confounders could explain the correlation of hydrophobicity with the compaction of unfolded states of foldable proteins.^24^ A key candidate for such a confounder is aromatic residues, which we found to induce strong compaction of the linkers. A tyrosine fraction of 0.1 was thus sufficient to contract the linker to dimensions observed for unfolded, but foldable proteins.^24^

### The additivity of IDP compaction

Several properties affect chain compaction, but how do they add up? Both charge and proline residues expand the linker, however the combination of the two did not expand the linker more than charge alone. This demonstrates that not all contributions to chain compaction are additive. A likely explanation is that rigidity added by proline residues can be accommodated inside the ensemble expanded by charge-charge repulsion. The reason the net charge density is such a good predictor of the compaction of IDPs, may thus be that the strongest effect dominates in the absence of additivity. The most likely explanation for the different threshold for charge expansion is thus the presence of compensating attractive interactions. This suggests an additivity between some factors, but likely not all, and demonstrates that much remains to be uncovered about sequence-compaction relationship of IDPs.

### The value of synthetic IDPs

Our present understanding of the relationship between sequence-structure in IDPs is mainly based on the study of natural proteins. In contrast, we have studied synthetic linkers never seen in nature. The synthetic linkers allow tight control over the physical properties of the linker, which is crucial for hypothesis testing. Experiments on synthetic IDPs are thus a natural step for critically evaluating our understanding of IDPs. Synthetic DNA has removed the need for ingenious cloning strategies used previously,^43^ leaving protein preparation as the major bottleneck. For artificial proteins spanning a range of physical properties, this can however be a major challenge. We have for example not been able to make our linkers in isolation yet. The fusion protein used here serves both as solubility-tags and a reporter system operating at nM concentrations. These are likely the key factors that have allowed us to study a broader range of IDPs than previous studies, and suggest that such fusion proteins provide useful tools for future investigations of sequence-structure relationships in IDPs.

## Acknowledgments

This work was supported by grants to M.K. from the “Young Investigator Program” of the Villum Foundation and the AIAS COFUND program (Agreement No. 609033). We wish to thank Birthe B. Kragelund, Mateusz Dyla and Xavier Warnet for critical comments to this manuscript, and Tanja Klymchuk for technical assistance.

## References

(1) Langeberg, L. K., and Scott, J. D. (2015) Signalling scaffolds and local organization of cellular behaviour. Nat. Rev. Mol. Cell Biol. 16, 232–244.

(2) Nussinov, R., Ma, B., and Tsai, C. J. (2013) A broad view of scaffolding suggests that scaffolding proteins can actively control regulation and signaling of multienzyme complexes through allostery. Biochim. Biophys. Acta - Proteins Proteomics 1834, 820–829.

(3) Good, M. C., Zalatan, J. G., and Lim, W. A. (2011) Scaffold Proteins: Hubs for Controlling the Flow of Cellular Information. Science (80-.). 332, 680–686.

(4) Dyson, H. J., and Wright, P. E. (2005) Intrinsically unstructured proteins and their functions. Nat. Rev. Mol. Cell Biol. 6, 197–208.

(5) Kitazawa, T., Igawa, T., Sampei, Z., Muto, A., Kojima, T., Soeda, T., Yoshihashi, K., Okuyama-Nishida, Y., Saito, H., Tsunoda, H., Suzuki, T., Adachi, H., Miyazaki, T., Ishii, S., Kamata-Sakurai, M., Iida, T., Harada, A., Esaki, K., Funaki, M., Moriyama, C., Tanaka, E., Kikuchi, Y., Wakabayashi, T., Wada, M., Goto, M., Toyoda, T., Ueyama, A., Suzuki, S., Haraya, K., Tachibana, T., Kawabe, Y., Shima, M., Yoshioka, A., and Hattori, K. (2012) A bispecific antibody to factors IXa and X restores factor VIII hemostatic activity in a hemophilia A model. Nat. Med. 18, 1570–1574.

(6) Greenwald, E. C., M. Redden J., Dodge-Kafka, K. L., and Saucerman, J. J. (2014) Scaffold state switching amplifies, accelerates, and insulates protein kinase c signaling. J. Biol. Chem. 289, 2353–2360.

(7) Cortese, M. S., Uversky, V. N., and Keith Dunker, A. (2008) Intrinsic disorder in scaffold proteins: Getting more from less. Prog. Biophys. Mol. Biol. 98, 85–106.

(8) Papaleo, E., Saladino, G., Lambrughi, M., Lindorff-Larsen, K., Gervasio, F. L., and Nussinov, R. (2016) The Role of Protein Loops and Linkers in Conformational Dynamics and Allostery. Chem. Rev. acs.chemrev.5b00623.

(9) Tompa, P. (2014) Multisteric Regulation by Structural Disorder in Modular Signaling Proteins: An Extension of the Concept of Allostery. Chem. Rev. 114, 6715–6732.

(10) Page, M. I., and Jencks, W. P. (1971) Entropic Contributions to Rate Accelerations in Enzymic and Intramolecular Reactions and the Chelate Effect. Proc. Natl. Acad. Sci. 68, 1678–1683.

(11) Krishnamurthy, V. M., Semetey, V., Bracher, P. J., Shen, N., and Whitesides, G. M. (2007) Dependence of effective molarity on linker length for an intramolecular protein-ligand system. J. Am. Chem. Soc. 129, 1312–1320.

(12) Gargano, J. M., Ngo, T., Kim, J. Y., Acheson, D. W. K., and Lees, W. J. (2001) Multivalent Inhibition of AB 5 Toxins. J. Am. Chem. Soc. 123, 12909–12910.

(13) Li, M., Cao, H., Lai, L., and Liu, Z. (2018) Disordered linkers in multidomain allosteric proteins: Entropic effect to favor the open state or enhanced local concentration to favor the closed state? Protein Sci. 27, 1600–1610.

(14) Mack, E. T., Snyder, P. W., Perez-Castillejos, R., Bilgi??er, B., Moustakas, D. T., Butte, M. J., and Whitesides, G. M. (2012) Dependence of avidity on linker length for a bivalent ligand-bivalent receptor model system. J. Am. Chem. Soc. 134, 333–345.

(15) Zhou, H. X. (2003) Quantitative account of the enhanced affinity of two linked scFvs specific for different epitopes on the same antigen. J. Mol. Biol. 329, 1–8.

(16) Timpe, L. C., and Peller, L. (1995) A random flight chain model for the tether of the Shaker K+ channel inactivation domain. Biophys. J. 69, 2415–2418.

(17) Diestler, D. J., and Knapp, E. W. (2010) Statistical Mechanics of the Stability of Multivalent Ligand - Receptor Complexes. J. Phys. Chem. C 114, 5287–5304.

(18) Diestler, D. J., and Knapp, E. W. (2008) Statistical mechanics of the stability of multivalent ligand-receptor complexes. Phys. Rev. Lett. 100, 178101.

(19) Borcherds, W., Becker, A., Chen, L., Chen, J., Chemes, L. B., and Daughdrill, G. W. (2017) Optimal Affinity Enhancement by a Conserved Flexible Linker Controls p53 Mimicry in MdmX. Biophys. J. 2038–2042.

(20) Sherry, K. P., Johnson, S. E., Hatem, C. L., Majumdar, A., and Barrick, D. (2015) Effects of Linker Length and Transient Secondary Structure Elements in the Intrinsically Disordered Notch RAM Region on Notch Signaling. J. Mol. Biol. 427, 3587–3597.

(21) Marsh, J. A., and Forman-Kay, J. D. (2010) Sequence determinants of compaction in intrinsically disordered proteins. Biophys. J. 98, 2374–2382.

(22) Bernadó, P., and Svergun, D. I. (2012) Analysis of intrinsically disordered proteins by small-angle X-ray scattering. Mol. Biosyst. 8, 151–167.

(23) Fuertes, G., Banterle, N., Ruff, K. M., Chowdhury, A., Mercadante, D., Koehler, C., Kachala, M., Estrada Girona, G., Milles, S., Mishra, A., Onck, P. R., Gräter, F., Esteban-Martín, S., Pappu, R. V., Svergun, D. I., and Lemke, E. A. (2017) Decoupling of size and shape fluctuations in heteropolymeric sequences reconciles discrepancies in SAXS vs. FRET measurements. Proc. Natl. Acad. Sci. 201704692.

(24) Hofmann, H., Soranno, a., Borgia, a., Gast, K., Nettels, D., and Schuler, B. (2012) Polymer scaling laws of unfolded and intrinsically disordered proteins quantified with single-molecule spectroscopy. Proc. Natl. Acad. Sci. 109, 16155–16160.

(25) Mao, a. H., Crick, S. L., Vitalis, a., Chicoine, C. L., and Pappu, R. V. (2010) Net charge per residue modulates conformational ensembles of intrinsically disordered proteins. Proc. Natl. Acad. Sci. 107, 8183–8188.

(26) Müller-Späth, S., Soranno, A., Hirschfeld, V., Hofmann, H., and Rüegger, S. (2010) Charge interactions can dominate the dimensions of intrinsically disordered proteins. Proc. Natl. Acad. Sci. 107, 14609–14614.

(27) Schuler, B., Soranno, A., Hofmann, H., and Nettels, D. (2016) Single-Molecule FRET Spectroscopy and the Polymer Physics of Unfolded and Intrinsically Disordered Proteins. Annu. Rev. Biophys. 45, 207–231.

(28) Das, R. K., Ruff, K. M., and Pappu, R. V. (2015) Relating sequence encoded information to form and function of intrinsically disordered proteins. Curr. Opin. Struct. Biol. 32, 102–112.

(29) Das, R. K., and Pappu, R. V. (2013) Conformations of intrinsically disordered proteins are influenced by linear sequence distributions of oppositely charged residues. Proc. Natl. Acad. Sci. U. S. A. 110, 13392–7.

(30) Martin, E. W., Holehouse, A. S., Grace, C. R., Hughes, A., Pappu, R. V, and Mittag, T. (2016) Sequence Determinants of the Conformational Properties of an Intrinsically Disordered Protein Prior to and upon Multisite Phosphorylation. J. Am. Chem. Soc. 138, 15323–15335.

(31) Studier, F. W. (2005) Protein production by auto-induction in high-density shaking cultures. Protein Expr. Purif. 41, 207–234.

(32) Kalthoff, C. (2003) A novel strategy for the purification of recombinantly expressed unstructured protein domains. J. Chromatogr. B Anal. Technol. Biomed. Life Sci. 786, 247–254.

(33) Zhang, J., Ma, Y., Taylor, S. S., and Tsien, R. Y. (2001) Genetically encoded reporters of protein kinase A activity reveal impact of substrate tethering. Proc. Natl. Acad. Sci. U. S. A. 98, 14997–15002.

(34) Bajar, B. T., Wang, E. S., Lam, A. J., Kim, B. B., Jacobs, C. L., Howe, E. S., Davidson, M. W., Lin, M. Z., and Chu, J. (2016) Improving brightness and photostability of green and red fluorescent proteins for live cell imaging and FRET reporting. Sci. Rep. 6, 1–12.

(35) Gnanapragasam, M. N., Scarsdale, J. N., Amaya, M. L., Webb, H. D., Desai, M. a, Walavalkar, N. M., Wang, S. Z., Zu Zhu, S., Ginder, G. D., and Williams, D. C. (2011) p66Alpha-MBD2 coiled-coil interaction and recruitment of Mi-2 are critical for globin gene silencing by the MBD2-NuRD complex. Proc. Natl. Acad. Sci. U. S. A. 108, 7487–92.

(36) Holehouse, A. S., Garai, K., Lyle, N., Vitalis, A., and Pappu, R. V. (2015) Quantitative assessments of the distinct contributions of polypeptide backbone amides versus side chain groups to chain expansion via chemical denaturation. J. Am. Chem. Soc. 137, 2984–2995.

(37) Kohn, J. E., Millett, I. S., Jacob, J., Zagrovic, B., Dillon, T. M., Cingel, N., Dothager, R. S., Seifert, S., Thiyagarajan, P., Sosnick, T. R., Hasan, M. Z., Pande, V. S., Ruczinski, I., Doniach, S., and Plaxco, K. W. (2004) Random-coil behavior and the dimensions of chemically unfolded proteins. Proc. Natl. Acad. Sci. 101, 12491–12496.

(38) Muller-Spath, S., Soranno, A., Hirschfeld, V., Hofmann, H., Ruegger, S., Reymond, L., Nettels, D., and Schuler, B. (2010) Charge interactions can dominate the dimensions of intrinsically disordered proteins. Proc. Natl. Acad. Sci. 107, 14609–14614.

(39) Wimley, W. C., and White, S. H. (1996) Experimentally determined hydrophobicity scales for membrane proteins. Nat. Struct. Biol. 3, 842–848.

(40) Wang, J., Choi, J. M., Holehouse, A. S., Lee, H. O., Zhang, X., Jahnel, M., Maharana, S., Lemaitre, R., Pozniakovsky, A., Drechsel, D., Poser, I., Pappu, R. V, Alberti, S., and Hyman, A. A. (2018) A Molecular Grammar Governing the Driving Forces for Phase Separation of Prion-like RNA Binding Proteins. Cell.

(41) Dobrynin, A. V. (2008) Theory and simulations of charged polymers: From solution properties to polymeric nanomaterials. Curr. Opin. Colloid Interface Sci. 13, 376–388.

(42) Borgia, A., Borgia, M. B., Bugge, K., Kissling, V. M., Heidarsson, P. O., Fernandes, C. B., Sottini, A., Soranno, A., Buholzer, K. J., Nettels, D., Kragelund, B. B., Best, R. B., and Schuler, B. (2018) Extreme disorder in an ultrahigh-affinity protein complex. Nature 555, 61–66.

(43) Evers, T. H., Faesen, A. C., Meijer, E. W., and Merkx, M. (2006) Quantitative Understanding of Energy Transfer between Fluorescent Proteins Connected via Flexible Peptide Linkers. Biochemistry 45, 13183–13192.

