## Supplemental information for "Effective concentrations enforced by intrinsically disordered linkers are governed by polymer physics"

Note: This is a pre-print, and has not yet been peer-reviewed! Copyright by the authors.

Note: This is a pre-print, and has not yet been peer-reviewed! Copyright by the authors.

### Protein sequences of constructs used:

#### Color code for features:

SA-Strep tag
mClover3
mRuby3
Unique restriction sites

#### Interaction partners:

MBD2
p66α

#### Fusion proteins:

MGSSHHHHHH SSGLVPRGSH MVSKGEELFT GVVPILVELD GDVNGHKFSV RGEGEGDATN
GKLTLKFICT TGKLPVPWPT LVTTFGYGVA CFSRYPDHMK QHDFFKSAMP EGYVQERTIS
FKDDGTYKTR AEVKFEGDTL VNRIELKGID FKEDGNILGH KLEYNFNSHY VYITADKQKN
CIKANFKIRH NVEDGSVQLA DHYQQNTPIG DGPVLLPDNH YLSHQSKLSK DPNEKRDHMV
LLEFVTAALE SGGEDPMVST GQSQSQSQSQ SVTDEDIRKQ EERAQQVRKK LEEALMADAS
(Variable linker)
GTPEERERMI KQLKEELRLE EAKLVLLKKL RQSTSQSQSQ SQSQSMVSKG EELIKENMRM
KVVMEGSVNG HQFKCTGEGE GRPYEGVQTM RIKVIEGGPL PFAFDILATS FMYGSRTFIK
YPADIPDFFK QSFPEGFTWE RVTRYEDGGV VTVTQDTSLE DGELVYNVKV RGVNFPSNGP
VMQKKTKGWE PNTEMMYPAD GGLRGYTDIA LKVDGGGHLH CNFVTTYRSK KTVGNIKMPG
VHAVDHRLER IEESDNETYV VQREVAVAKY SNLGGGMDEL YKQSQSQSWS HPQFEK

#### MBD2 WT peptide:

MGSSHHHHHH SSGLVPRGSH MQSQSQSQSQ S VTDEDIRKQ EERVQQVRKK LEEALMADAS
GSGSGSGSGS Y

Note: This is a pre-print, and has not yet been peer-reviewed! Copyright by the authors.

#### Supplementary figures:

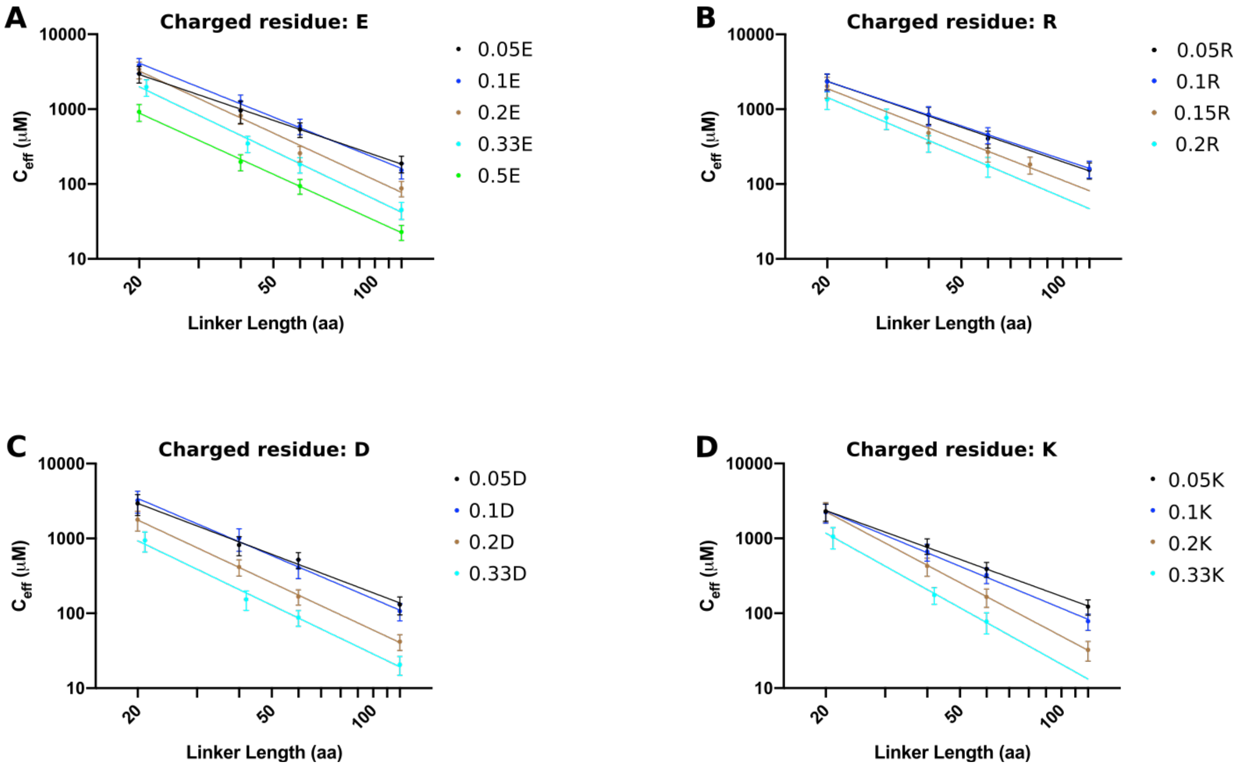


Fig. S1: Power law fits to experimentally determined effective concentrations for linkers containing different types of charged residues

Note: This is a pre-print, and has not yet been peer-reviewed! Copyright by the authors.

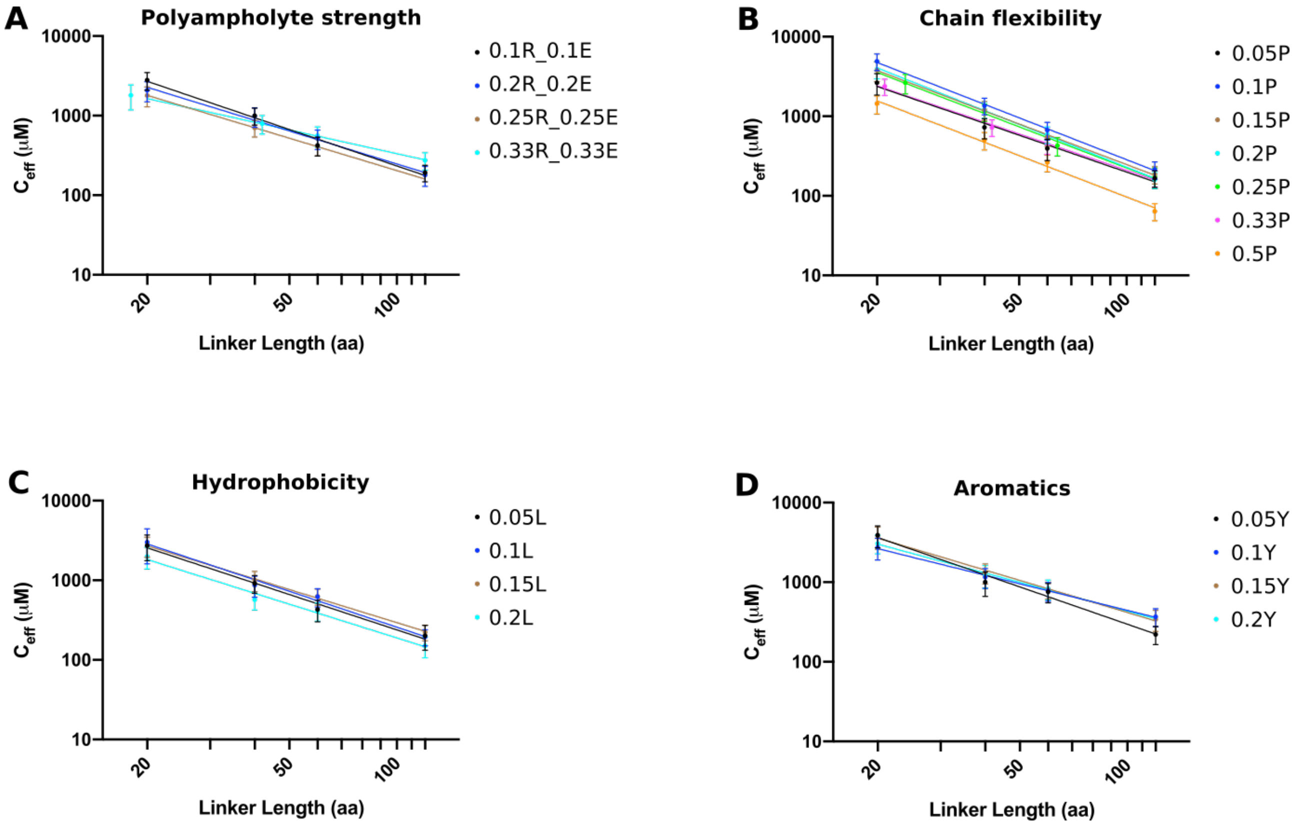


**Fig. S2:** Power law fits to experimentally determined effective concentrations for linkers with variations in polyampholyte strength, chain flexibility, hydrophobicity and aromatic residue content.

Note: This is a pre-print, and has not yet been peer-reviewed! Copyright by the authors.

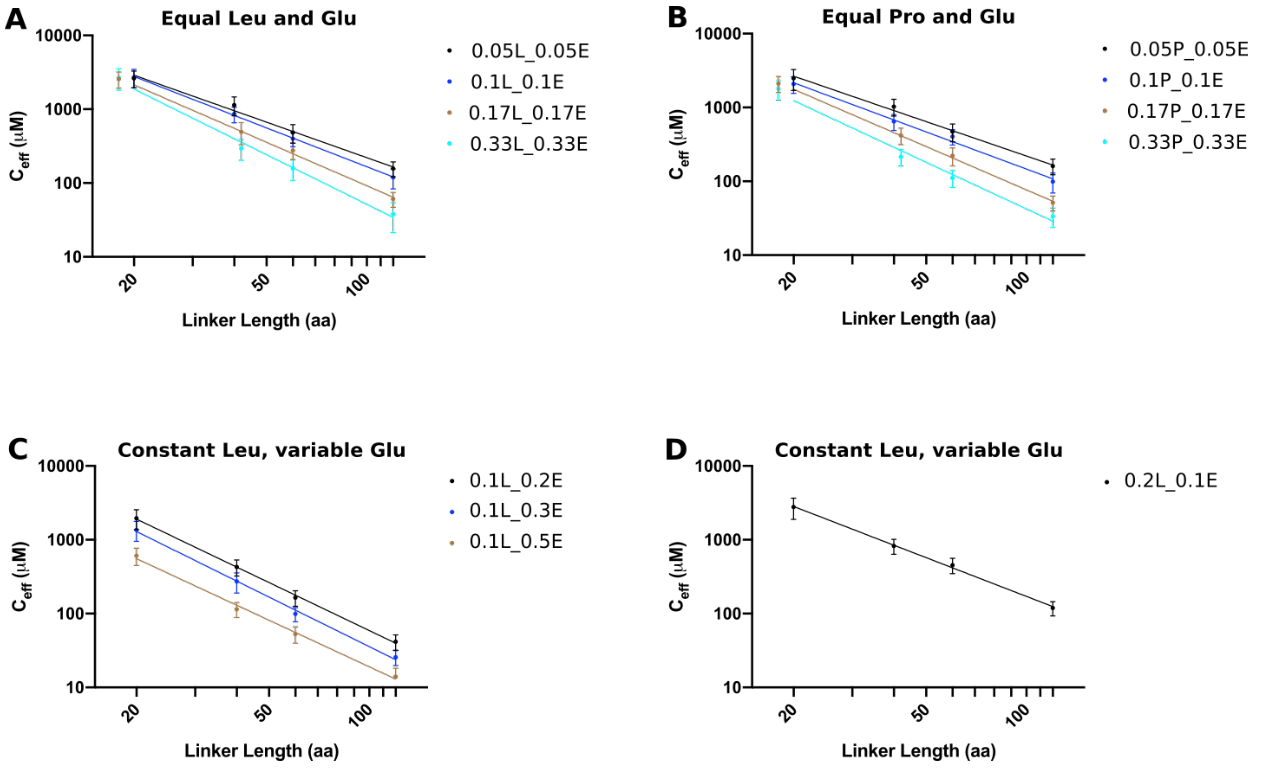


**Fig. S3:** Power law fits to experimentally determined effective concentrations for linkers with combinations of charged residues and leucine or proline.

Note: This is a pre-print, and has not yet been peer-reviewed! Copyright by the authors.

#### Table S1: Full sequence of all linkers used:

| Linker name | Linker sequence |
|---|---|
| **GS** | |
| $GS_{20}$ | GSGSGSGSGSGSGSGSGSGS |
| $GS_{30}$ | GSGSGSGSGSGSGSGSGSGSGSGSGSGSGS |
| $GS_{40}$ | GSGSGSGSGSGSGSGSGSGSGSGSGSGSGSGSGSGSGSGS |
| $GS_{60}$ | GSGSGSGSGSGSGSGSGSGSGSGSGSGSGSGSGSGSGSGSGSGSGSGSGSGSGSGSGSGS |
| $GS_{120}$ | GSGSGSGSGSGSGSGSGSGSGSGSGSGSGSGSGSGSGSGSGSGSGSGSGSGSGSGSGSGSGSGSGSGSGSGSGSGS<br>GSGSGSGSGSGSGSGSGSGSGSGSGSGSGSGSGSGSGSGSGSGS |
| **E** | |
| $0.05E_{20}$ | GSGSGSGSGESGSGSGSGSG |
| $0.05E_{40}$ | GSGSGSGSGESGSGSGSGSGSGSGSGSGSGESGSGSGSGS |
| $0.05E_{60}$ | GSGSGSGSGESGSGSGSGSGSGSGSGSGSGESGSGSGSGSGSGSGSGSGSGESGSGSGSG |
| $0.05E_{120}$ | GSGSGSGSGESGSGSGSGSGSGSGSGSGSGESGSGSGSGSGSGSGSGSGSGESGSGSGSGSGSGSGSGSGSGESGSG<br>SGSGSGSGSGESGSGSGSGSGSGSGSGSGSGESG |
| $0.1E_{20}$ | SGSGSGSGS |
| $0.1E_{40}$ | GSGSEGSGSGSGSGESGSGSGSGSEGSGSGSGSGESGSGS |
| $0.1E_{60}$ | GSGSEGSGSGSGSGESGSGSGSGSEGSGSGSGSGESGSGSGSGSEGSGSGSGSGESGSGS |
| $0.1E_{120}$ | GSGSEGSGSGSGSGESGSGSGSGSEGSGSGSGSGESGSGSGSGSEGSGSGSGSGESGSGSGSGSEGSGSGSGSGESG<br>SGSGSGSEGSGSGSGSGESGSGSGSGSEGSGSGSGSGESGSGS |
| $0.2E_{20}$ | GSEGSGSEGSGSEGSGSEGS |
| $0.2E_{40}$ | GSEGSGSEGSGSEGSGSEGSGSEGSGSEGSGSEGSGSEGS |
| $0.2E_{60}$ | GSEGSGSEGSGSEGSGSEGSGSEGSGSEGSGSEGSGSEGSGSEGSGSEGSGSEGSGSEGS |
| $0.2E_{120}$ | GSEGSGSEGSGSEGSGSEGSGSEGSGSEGSGSEGSGSEGSGSEGSGSEGSGSEGSGSEGSGSEGSGSEGSGSEGSGSE<br>GSGSEGSGSEGSGSEGSGSEGSGSEGSGSEGSGSEGSGSEGS |
| $0.33E_{21}$ | GESGESGESGESGESGESGES |
| $0.33E_{42}$ | GESGESGESGESGESGESGESGESGESGESGESGESGESGES |
| $0.33E_{60}$ | GESGESGESGESGESGESGESGESGESGESGESGESGESGESGESGESGESGESGESGES |
| $0.33E_{120}$ | GESGESGESGESGESGESGESGESGESGESGESGESGESGESGESGESGESGESGESGESGESGESGESGESGESGESG<br>ESGESGESGESGESGESGESGESGESGESGESGESGESGES |
| $0.5E_{20}$ | GESEGESEGESEGESEGESE |
| $0.5E_{40}$ | GESEGESEGESEGESEGESEGESEGESEGESEGESEGESE |
| $0.5E_{60}$ | GESEGESEGESEGESEGESEGESEGESEGESEGESEGESEGESEGESEGESEGESEGESE |
| $0.5E_{120}$ | GESEGESEGESEGESEGESEGESEGESEGESEGESEGESEGESEGESEGESEGESEGESEGESEGESEGESEGESEG<br>ESEGESEGESEGESEGESEGESEGESEGESEGESEGESE |
| **R** | |
| $0.05R_{20}$ | GSGSGSGSGRSGSGSGSGSG |
| $0.05R_{40}$ | GSGSGSGSGRSGSGSGSGSGSGSGSGSGSGRSGSGSGSGS |
| $0.05R_{60}$ | GSGSGSGSGRSGSGSGSGSGSGSGSGSGSGRSGSGSGSGSGSGSGSGSGSGRSGSGSGSG |
| $0.05R_{120}$ | GSGSGSGSGRSGSGSGSGSGSGSGSGSGSGRSGSGSGSGSGSGSGSGSGSGRSGSGSGSGSGSGSGSGSGSGRSGSG<br>SGSGSGSGSGRSGSGSGSGSGSGSGSGSGSGRSGSGSGSGSGS |
| $0.1R_{20}$ | GSGSRGSGSGSGSGRSGSGS |
| $0.1R_{40}$ | GSGSRGSGSGSGSGRSGSGSGSGSRGSGSGSGSGRSGSGS |
| $0.1R_{60}$ | GSGSRGSGSGSGSGRSGSGSGSGSRGSGSGSGSGRSGSGSGSGSRGSGSGSGSGRSGSGS |
| $0.1R_{120}$ | GSGSRGSGSGSGSGRSGSGSGSGSRGSGSGSGSGRSGSGSGSGSRGSGSGSGSGRSGSGSGSGSRGSGSGSGSGRSG<br>SGSGSGSRGSGSGSGSGRSGSGSGSGSRGSGSGSGSGRSGSGS |

| | |
|---|---|
| $0.15R_{20}$ | GSGRSGSGSRGSGSGSRGSG |
| $0.15R_{40}$ | GSGRSGSGSRGSGSGSRGSGSGSRGSGSGSRGSGSGRSGS |
| $0.15R_{60}$ | GSGRSGSGSRGSGSGSRGSGSGSRGSGSGSRGSGSGRSGSGSGRSGSGSRGSGSGSRGSG |
| $0.15R_{80}$ | GSGSGRSGSGSRGSGSGSRGSGSGRSGSGSRGSGSGRSGSGSGRSGSGSRGSGSGRSGSGSRGSGSGSRGSGSGRSG SGS |
| $0.2R_{20}$ | GSRGSGSRGSGSRGSGSRGS |
| $0.2R_{30}$ | GSGRSGSGRSGSRGSGSRGSGRSGSGRSGS |
| $0.2R_{40}$ | GSRGSGSRGSGSRGSGSRGSGSRGSGSRGSGSRGSGSRGS |
| $0.2R_{60}$ | GSRGSGSRGSGSRGSGSRGSGSRGSGSRGSGSRGSGSRGSGSRGSGSRGSGSRGSGSRGS |
| **L** | |
| $0.05L_{20}$ | GSGSGSGSGLSGSGSGSGSG |
| $0.05L_{40}$ | GSGSGSGSGLSGSGSGSGSGSGSGSGSGLSGSGSGSGS |
| $0.05L_{60}$ | GSGSGSGSGLSGSGSGSGSGSGSGSGSGLSGSGSGSGSGSGSGSGLSGSGSGSGSG |
| $0.05L_{120}$ | GSGSGSGSGLSGSGSGSGSGSGSGSGSGLSGSGSGSGSGSGSGSGLSGSGSGSGSGSGSGSGLSGSGSGSGS GSGSGSGSGSGLSGSGSGSGSGSGSGSGLSGSGSGSGSGS |
| $0.1L_{20}$ | GSGSLGSGSGSGSGLSGSGS |
| $0.1L_{40}$ | GSGSLGSGSGSGSGLSGSGSGSGSLGSGSGSGSGLSGSGS |
| $0.1L_{60}$ | GSGSLGSGSGSGSGLSGSGSGSGSLGSGSGSGSGLSGSGSGSGSLGSGSGSGSGLSGSGS |
| $0.1L_{120}$ | GSGSLGSGSGSGSGLSGSGSGSGSLGSGSGSGSGLSGSGSGSGSLGSGSGSGSGLSGSGSGSGSLGSGSGSGSGLSGS GSGSGSLGSGSGSGSGLSGSGSGSGSLGSGSGSGSGLSGSGS |
| $0.15L_{20}$ | GSGLSGSGSLGSGSGSLGSG |
| $0.15L_{40}$ | GSGLSGSGSLGSGSGSLGSGSGSLGSGSGSLGSGSGLSGS |
| $0.15L_{60}$ | GSGLSGSGSLGSGSGSLGSGSGSLGSGSGSLGSGSGLSGSGSGLSGSGSLGSGSGSLGSG |
| $0.15L_{120}$ | GSGLSGSGSLGSGSGSLGSGSGSLGSGSGSLGSGSGLSGSGSGLSGSGSLGSGSGSLGSGSGLSGSGSLGSGSGSLGSG SGSLGSGSGSLGSGSGLSGSGSGLSGSGSLGSGSGSLGSGS |
| $0.2L_{20}$ | GSLGSGSLGSGSLGSGSLGS |
| $0.2L_{40}$ | GSLGSGSLGSGSLGSGSLGSGSLGSGSLGSGSLGSGSLGS |
| $0.2L_{60}$ | GSLGSGSLGSGSLGSGSLGSGSLGSGSLGSGSLGSGSLGSGSLGSGSLGSGSLGSGSLGS |
| $0.2L_{120}$ | GSLGSGSLGSGSLGSGSLGSGSLGSGSLGSGSLGSGSLGSGSLGSGSLGSGSLGSGSLGSGSLGSGSLGSGSLGSGSLGS GSLGSGSLGSGSLGSGSLGSGSLGSGSLGSGSLGSGSLGS |
| **P** | |
| $0.05P_{20}$ | GSGSGSGSGPSGSGSGSGSG |
| $0.05P_{40}$ | GSGSGSGSGPSGSGSGSGSGSGSGSGSGPSGSGSGSGS |
| $0.05P_{60}$ | GSGSGSGSGPSGSGSGSGSGSGSGSGSGPSGSGSGSGSGSGSGSGPSGSGSGSGSG |
| $0.05P_{120}$ | GSGSGSGSGPSGSGSGSGSGSGSGSGSGPSGSGSGSGSGSGSGSGPSGSGSGSGSGSGSGSGPSGSGSGSG SGSGSGSGSGSGPSGSGSGSGSGSGSGSGPSGSGSGSGSGS |
| $0.1P_{20}$ | GSGSPGSGSGSGSGPSGSGS |
| $0.1P_{40}$ | GSGSPGSGSGSGSGPSGSGSGSGSPGSGSGSGSGPSGSGS |
| $0.1P_{60}$ | GSGSPGSGSGSGSGPSGSGSGSGSPGSGSGSGSGPSGSGSGSGSPGSGSGSGSGPSGSGS |
| $0.1P_{120}$ | GSGSPGSGSGSGSGPSGSGSGSGSPGSGSGSGSGPSGSGSGSGSPGSGSGSGSGPSGSGSGSGSPGSGSGSGSGPSG SGSGSGSPGSGSGSGSGPSGSGSGSGSPGSGSGSGSGPSGSGS |
| $0.15P_{20}$ | GSGPSGSGSPGSGSGSPGSG |
| $0.15P_{40}$ | GSGPSGSGSPGSGSGSPGSGSGSPGSGSGSPGSGSGPSGS |
| $0.15P_{60}$ | GSGPSGSGSPGSGSGSPGSGSGSPGSGSGSPGSGSGPSGSGSGPSGSGSPGSGSGSPGSG |
| $0.15P_{120}$ | GSGPSGSGSPGSGSGSPGSGSGSPGSGSGSPGSGSGPSGSGSGPSGSGSPGSGSGSPGSGSGPSGSGSPGSGSGSPG SGSGSPGSGSGSPGSGSGPSGSGSGPSGSGSPGSGSGSPGSGS |

Note: This is a pre-print, and has not yet been peer-reviewed! Copyright by the authors.

| | |
|---|---|
| $0.2P_{20}$ | GSPGSGSPGSGSPGSGSPGS |
| $0.2P_{40}$ | GSPGSGSPGSGSPGSGSPGSGSPGSGSPGSGSPGSGSPGS |
| $0.2P_{60}$ | GSPGSGSPGSGSPGSGSPGSGSPGSGSPGSGSPGSGSPGSGSPGSGSPGSGSPGSGSPGS |
| $0.2P_{120}$ | GSPGSGSPGSGSPGSGSPGSGSPGSGSPGSGSPGSGSPGSGSPGSGSPGSGSPGSGSPGSGSPGSGSPGSGSPGSGSPGSGSPGSGSPGSGSPGSGSPGSGSPGSGSPGSGSPGSGSPGS |
| $0.25P_{24}$ | GSPGSGPSGSPGSPGSGPSGSPGS |
| $0.25P_{40}$ | GSPGSGPSGSPGSGPSGSPGSPGSGPSGSPGSGPSGSPGS |
| $0.25P_{64}$ | GSPGSGPSGSPGSGPSGSPGSGPSGSPGSGPSGPSGSPGSGPSGSPGSGPSGSPGSGPSGSPGS |
| $0.25P_{120}$ | GSPGSGPSGSPGSGPSGSPGSGPSGSPGSGPSGSPGSGPSGSPGSGPSGSPGSGPSGSPGSPGSGPSGSPGSGPSGSPGSGPSGSPGSGPSGSPGSGPSGSPGSGPSGSPGSGPSGSPGS |
| $0.33P_{21}$ | GPSGPSGPSGPSGPSGPSGPS |
| $0.33P_{42}$ | GPSGPSGPSGPSGPSGPSGPSGPSGPSGPSGPSGPSGPSGPS |
| $0.33P_{60}$ | GPSGPSGPSGPSGPSGPSGPSGPSGPSGPSGPSGPSGPSGPSGPSGPSGPSGPSGPSGPS |
| $0.33P_{120}$ | GPSGPSGPSGPSGPSGPSGPSGPSGPSGPSGPSGPSGPSGPSGPSGPSGPSGPSGPSGPSGPSGPSGPSGPSGPSGPSGPSGPSGPSGPSGPSGPSGPSGPSGPSGPSGPSGPSGPSGPS |
| $0.5P_{20}$ | GPSPGPSPGPSPGPSPGPSP |
| $0.5P_{40}$ | GPSPGPSPGPSPGPSPGPSPGPSPGPSPGPSPGPSPGPSP |
| $0.5P_{60}$ | GPSPGPSPGPSPGPSPGPSPGPSPGPSPGPSPGPSPGPSPGPSPGPSPGPSPGPSPGPSP |
| $0.5P_{120}$ | GPSPGPSPGPSPGPSPGPSPGPSPGPSPGPSPGPSPGPSPGPSPGPSPGPSPGPSPGPSPGPSPGPSPGPSPGPSPGPSPGPSPGPSPGPSPGPSPGPSPGPSPGPSPGPSPGPSPGPSP |
| **PE** | |
| $0.05PE_{20}$ | GSGSPGSGSGSGSGESGSGS |
| $0.05PE_{40}$ | GSGSPGSGSGSGSGESGSGSGSGSPGSGSGSGSGESGSGS |
| $0.05PE_{60}$ | GSGSPGSGSGSGSGESGSGSGSGSPGSGSGSGSGESGSGSGSGSPGSGSGSGSGESGSGS |
| $0.05PE_{120}$ | GSGSPGSGSGSGSGESGSGSGSGSPGSGSGSGSGESGSGSGSGSPGSGSGSGSGESGSGSGSGSPGSGSGSGSGESGSGSGSGSPGSGSGSGSGESGSGSGSGSPGSGSGSGSGESGSGS |
| $0.1PE_{20}$ | GSPGSGSEGSGSPGSGSEGS |
| $0.1PE_{40}$ | GSPGSGSEGSGSPGSGSEGSGSPGSGSEGSGSPGSGSEGS |
| $0.1PE_{60}$ | GSPGSGSEGSGSPGSGSEGSGSPGSGSEGSGSPGSGSEGSGSPGSGSEGSGSPGSGSEGS |
| $0.1PE_{120}$ | GSPGSGSEGSGSPGSGSEGSGSPGSGSEGSGSPGSGSEGSGSPGSGSEGSGSPGSGSEGSGSPGSGSEGSGSPGSGSEGSGSPGSGSEGSGSPGSGSEGSGSPGSGSEGSGSPGSGSEGS |
| $0.167PE_{18}$ | GPSGESGPSGESGPSGES |
| $0.167PE_{42}$ | GPSGESGPSGESGPSGESGPSGESGPSGESGPSGESGPSGES |
| $0.167PE_{60}$ | GPSGESGPSGESGPSGESGPSGESGPSGESGPSGESGPSGESGPSGESGPSGESGPSGES |
| $0.167PE_{120}$ | GPSGESGPSGESGPSGESGPSGESGPSGESGPSGESGPSGESGPSGESGPSGESGPSGESGPSGESGPSGESGPSGESGPSGESGPSGESGPSGESGPSGESGPSGESGPSGESGPSGES |
| $0.33PE_{18}$ | GPESPEGPESPEGPESPE |
| $0.33PE_{42}$ | GPESPEGPESPEGPESPEGPESPEGPESPEGPESPEGPESPE |
| $0.33PE_{60}$ | GPESPEGPESPEGPESPEGPESPEGPESPEGPESPEGPESPEGPESPEGPESPEGPESPE |
| $0.33PE_{120}$ | GPESPEGPESPEGPESPEGPESPEGPESPEGPESPEGPESPEGPESPEGPESPEGPESPEGPESPEGPESPEGPESPEGPESPEGPESPEGPESPEGPESPEGPESPEGPESPEGPESPE |
| **LE** | |
| $0.05LE_{20}$ | GSGSLGSGSGSGSGESGSGS |
| $0.05LE_{40}$ | GSGSLGSGSGSGSGESGSGSGSGSLGSGSGSGSGESGSGS |
| $0.05LE_{60}$ | GSGSLGSGSGSGSGESGSGSGSGSLGSGSGSGSGESGSGSGSGSLGSGSGSGSGESGSGS |
| $0.05LE_{120}$ | GSGSLGSGSGSGSGESGSGSGSGSLGSGSGSGSGESGSGSGSGSLGSGSGSGSGESGSGSGSGSLGSGSGSGSGESGSGSGSGSLGSGSGSGSGESGSGSGSGSLGSGSGSGSGESGSGS |

Note: This is a pre-print, and has not yet been peer-reviewed! Copyright by the authors.

| | |
|---|---|
| 0.1LE$_{20}$ | GSLGSGSEGSGSLGSGSEGS |
| 0.1LE$_{40}$ | GSLGSGSEGSGSLGSGSEGSGSLGSGSEGSGSLGSGSEGS |
| 0.1LE$_{60}$ | GSLGSGSEGSGSLGSGSEGSGSLGSGSEGSGSLGSGSEGSGSLGSGSEGSGSLGSGSEGS |
| 0.1LE$_{120}$ | GSLGSGSEGSGSLGSGSEGSGSLGSGSEGSGSLGSGSEGSGSLGSGSEGSGSLGSGSEGSGSLGSGSEGSGSLGSGSEG SGSLGSGSEGSGSLGSGSEGSGSLGSGSEGSGSLGSGSEGS |
| 0.167LE$_{18}$ | GLSGESGLSGESGLSGES |
| 0.167LE$_{42}$ | GLSGESGLSGESGLSGESGLSGESGLSGESGLSGESGLSGES |
| 0.167LE$_{60}$ | GLSGESGLSGESGLSGESGLSGESGLSGESGLSGESGLSGESGLSGESGLSGESGLSGES |
| 0.167LE$_{120}$ | GLSGESGLSGESGLSGESGLSGESGLSGESGLSGESGLSGESGLSGESGLSGESGLSGESGLSGESGLSGESGLSGESGLS GESGLSGESGLSGESGLSGESGLSGESGLSGESGLSGES |
| 0.33LE$_{18}$ | GLESLEGLESLEGLESLE |
| 0.33LE$_{42}$ | GLESLEGLESLEGLESLEGLESLEGLESLEGLESLEGLESLE |
| 0.33LE$_{60}$ | GLESLEGLESLEGLESLEGLESLEGLESLEGLESLEGLESLEGLESLEGLESLEGLESLE |
| 0.33LE$_{120}$ | GLESLEGLESLEGLESLEGLESLEGLESLEGLESLEGLESLEGLESLEGLESLEGLESLEGLESLEGLESLEGLESLEGL ESLEGLESLEGLESLEGLESLEGLESLEGLESLE |
| **L+E** | |
| 0.1L_0.2E$_{20}$ | GSEGSLGSEGSEGSLGSEGS |
| 0.1L_0.2E$_{40}$ | GSGESGLSGESGESGLSGESGESGLSGESGESGLSGESGS |
| 0.1L_0.2E$_{60}$ | GSGESGLSGESGESGLSGESGSEGSLGSEGSEGSLGSEGSGESGLSGESGESGLSGESGS |
| 0.1L_0.2E$_{120}$ | GSGESGLSGESGESGSLGSEGSEGSLGSGESGESGLSGESGSEGSLGSEGSEGSGLSGESGESGLSGSEGSEGSLGSEGS GESGLSGESGESGSLGSEGSEGSLGSGESGESGLSGESGS |
| 0.1L_0.3E$_{20}$ | GSEGESLGESGESLGESEGS |
| 0.1L_0.3E$_{40}$ | GSEGESLGSEGESEGSLGESEGSLGESEGSEGLSEGSEGS |
| 0.1L_0.3E$_{60}$ | GSEGESGLSEGESGESLGSEGESEGSLGESGESLGSEGESEGSLGESGESEGLSGESEGS |
| 0.1L_0.3E$_{120}$ | GSEGESLGSEGESGESLGESGESEGSLGESGESEGSLGESGESEGSLGESGESEGSLGESGESLGSEGESGESLGSEGESG ESLGSEGESGESLGSEGESGESLGESGESEGSLGESEGS |
| 0.1L_0.5E$_{20}$ | GESEEGLESEEGELSEEGES |
| 0.1L_0.5E$_{40}$ | GESEEGESLEGEESEGLESEEGELSEGEESELGESEEGES |
| 0.1L_0.5E$_{60}$ | GESEGEESELGESEEGELSEGEESELGESEEGESLEGEESEGLESEEGESLEGEESEGES |
| 0.1L_0.5E$_{120}$ | GESEGEESEGLESEEGESELGESEEGESLEGEESEGELSEGEESEGLESEEGESELGESEEGESLEGESEEGELSEGEESEGL ESEGEESELGESEEGESLEGESEEGELSEGEESEGES |
| **E+L** | |
| 0.1E_0.2L$_{20}$ | GSLGSEGSLGSLGSEGSLGS |
| 0.1E_0.2L$_{40}$ | GSGLSGESGLSGLSGESGLSGLSGESGLSGLSGESGLSGS |
| 0.1E_0.2L$_{60}$ | GSGLSGESGLSGLSGESGLSGSLGSEGSLGSLGSEGSLGSGLSGESGLSGLSGESGLSGS |
| 0.1E_0.2L$_{120}$ | GSGLSGESGLSGLSGSEGSLGSLGSEGSGLSGLSGESGLSGSLGSEGSLGSLGSGESGLSGLSGESGSLGSLGSEGSLGSG LSGESGLSGLSGSEGSLGSLGSEGSGLSGLSGESGLSGS |
| **RE** | |
| 0.1RE$_{20}$ | GSRGSGSEGSGSRGSGSEGS |
| 0.1RE$_{40}$ | GSRGSGSEGSGSRGSGSEGSGSRGSGSEGSGSRGSGSEGS |
| 0.1RE$_{60}$ | GSRGSGSEGSGSRGSGSEGSGSRGSGSEGSGSRGSGSEGSGSRGSGSEGSGSRGSGSEGS |
| 0.1RE$_{120}$ | GSRGSGSEGSGSRGSGSEGSGSRGSGSEGSGSRGSGSEGSGSRGSGSEGSGSRGSGSEGSGSRGSGSEGSGSRGSGSE GSGSRGSGSEGSGSRGSGSEGSGSRGSGSEGSGSRGSGSEGS |
| 0.2RE$_{20}$ | GSRGESRGESGRSEGRSEGS |
| 0.2RE$_{40}$ | GRSEGSRGESGRSEGSRGESGRSEGSRGESGRSEGSRGES |
| 0.2RE$_{60}$ | GRSEGSRGESGRSEGSRGESGRSEGSRGESGRSEGSRGESGRSEGSRGESGRSEGSRGES |

| | |
|---|---|
| $0.2RE_{120}$ | GRSEGSRGESGRSEGSRGESGRSEGSRGESGRSEGSRGESGRSEGSRGESGRSEGSRGESGRSEGSRGESGRSEGSRGESGRSEGSRGESGRSEGSRGESGRSEGSRGESGRSEGSRGES |
| $0.25RE_{20}$ | GRSEGRSEGRSEGRSEGRSE |
| $0.25RE_{40}$ | GRSEGRSEGRSEGRSEGRSEGRSEGRSEGRSEGRSEGRSE |
| $0.25RE_{60}$ | GRSEGRSEGRSEGRSEGRSEGRSEGRSEGRSEGRSEGRSEGRSEGRSEGRSEGRSEGRSE |
| $0.33RE_{18}$ | GRESREGRESREGRESRE |
| $0.33RE_{42}$ | GRESREGRESREGRESREGRESREGRESREGRESREGRESRE |
| $0.33RE_{60}$ | GRESREGRESREGRESREGRESREGRESREGRESREGRESREGRESREGRESREGRESRE |
| $0.33RE_{120}$ | GRESREGRESREGRESREGRESREGRESREGRESREGRESREGRESREGRESREGRESREGRESREGRESREGRESREGRESREGRESREGRESREGRESREGRESREGRESREGRESRE |
| **D** | |
| $0.05D_{20}$ | GSGSGSGSGDSGSGSGSGSG |
| $0.05D_{40}$ | GSGSGSGSGDSGSGSGSGSGSGSGSGSGDSGSGSGSGS |
| $0.05D_{60}$ | GSGSGSGSGDSGSGSGSGSGSGSGSGSGDSGSGSGSGSGSGSGSGDSGSGSGSGSG |
| $0.05D_{120}$ | GSGSGSGSGDSGSGSGSGSGSGSGSGSGDSGSGSGSGSGSGSGSGDSGSGSGSGSGSGSGSGDSGSGSGSGSGSGSGSGSGSGDSGSGSGSGSGSGSGSGDSGSGSGSGSGS |
| $0.1D_{20}$ | GSGSDGSGSGSGSGDSGSGS |
| $0.1D_{40}$ | GSGSDGSGSGSGSGDSGSGSGSGSDGSGSGSGSGDSGSGS |
| $0.1D_{60}$ | GSGSDGSGSGSGSGDSGSGSGSGSDGSGSGSGSGDSGSGSGSGSDGSGSGSGSGDSGSGS |
| $0.1D_{120}$ | GSGSDGSGSGSGSGDSGSGSGSGSDGSGSGSGSGDSGSGSGSGSDGSGSGSGSGDSGSGSGSGSDGSGSGSGSGDSGSGSGSGSDGSGSGSGSGDSGSGSGSGSDGSGSGSGSGDSGSGS |
| $0.2D_{20}$ | GSDGSGSDGSGSDGSGSDGS |
| $0.2D_{40}$ | GSDGSGSDGSGSDGSGSDGSGSDGSGSDGSGSDGSGSDGS |
| $0.2D_{60}$ | GSDGSGSDGSGSDGSGSDGSGSDGSGSDGSGSDGSGSDGSGSDGSGSDGSGSDGSGSDGS |
| $0.2D_{120}$ | GSDGSGSDGSGSDGSGSDGSGSDGSGSDGSGSDGSGSDGSGSDGSGSDGSGSDGSGSDGSGSDGSGSDGSGSDGSGSDGSGSDGSGSDGSGSDGSGSDGSGSDGSGSDGSGSDGSGSDGS |
| $0.33D_{21}$ | GDSGDSGDSGDSGDSGDSGDS |
| $0.33D_{42}$ | GDSGDSGDSGDSGDSGDSGDSGDSGDSGDSGDSGDSGDSGDS |
| $0.33D_{60}$ | GDSGDSGDSGDSGDSGDSGDSGDSGDSGDSGDSGDSGDSGDSGDSGDSGDSGDSGDSGDS |
| $0.33D_{120}$ | GDSGDSGDSGDSGDSGDSGDSGDSGDSGDSGDSGDSGDSGDSGDSGDSGDSGDSGDSGDSGDSGDSGDSGDSGDSGDSGDSGDSGDSGDSGDSGDSGDSGDSGDSGDSGDSGDSGDSGDS |
| **K** | |
| $0.05K_{20}$ | GSGSGSGSGKSGSGSGSGSG |
| $0.05K_{40}$ | GSGSGSGSGKSGSGSGSGSGSGSGSGSGKSGSGSGSGS |
| $0.05K_{60}$ | GSGSGSGSGKSGSGSGSGSGSGSGSGSGKSGSGSGSGSGSGSGSGKSGSGSGSGSG |
| $0.05K_{120}$ | GSGSGSGSGKSGSGSGSGSGSGSGSGSGKSGSGSGSGSGSGSGSGKSGSGSGSGSGSGSGSGKSGSGSGSGSGSGSGSGSGSGKSGSGSGSGSGSGSGSGKSGSGSGSGSGS |
| $0.1K_{20}$ | GSGSKGSGSGSGSGKSGSGS |
| $0.1K_{40}$ | GSGSKGSGSGSGSGKSGSGSGSGSKGSGSGSGSGKSGSGS |
| $0.1K_{60}$ | GSGSKGSGSGSGSGKSGSGSGSGSKGSGSGSGSGKSGSGSGSGSKGSGSGSGSGKSGSGS |
| $0.1K_{120}$ | GSGSKGSGSGSGSGKSGSGSGSGSKGSGSGSGSGKSGSGSGSGSKGSGSGSGSGKSGSGSGSGSKGSGSGSGSGKSGSGSGSGSKGSGSGSGSGKSGSGSGSGSKGSGSGSGSGKSGSGS |
| $0.2K_{20}$ | GSKGSGSKGSGSKGSGSKGS |
| $0.2K_{40}$ | GSKGSGSKGSGSKGSGSKGSGSKGSGSKGSGSKGSGSKGS |
| $0.2K_{60}$ | GSKGSGSKGSGSKGSGSKGSGSKGSGSKGSGSKGSGSKGSGSKGSGSKGSGSKGSGSKGS |
| $0.2K_{120}$ | GSKGSGSKGSGSKGSGSKGSGSKGSGSKGSGSKGSGSKGSGSKGSGSKGSGSKGSGSKGSGSKGSGSKGSGSKGSGSKGSGSKGSGSKGSGSKGSGSKGSGSKGSGSKGSGSKGSGSKGS |

| | |
|---|---|
| 0.33$K_{21}$ | GKSGKSGKSGKSGKSGKSGKS |
| 0.33$K_{42}$ | GKSGKSGKSGKSGKSGKSGKSGKSGKSGKSGKSGKSGKSGKS |
| 0.33$K_{60}$ | GKSGKSGKSGKSGKSGKSGKSGKSGKSGKSGKSGKSGKSGKSGKSGKSGKSGKSGKSGKS |
| **Y** | |
| 0.05$Y_{20}$ | GSGSGSGSGYSGSGSGSGSG |
| 0.05$Y_{40}$ | GSGSGSGSGYSGSGSGSGSGSGSGSGSGYSGSGSGSGS |
| 0.05$Y_{60}$ | GSGSGSGSGYSGSGSGSGSGSGSGSGSGYSGSGSGSGSGSGSGSGYSGSGSGSGSG |
| 0.05$Y_{120}$ | GSGSGSGSGYSGSGSGSGSGSGSGSGSGYSGSGSGSGSGSGSGSGYSGSGSGSGSGSGSGSGYSGSGSGSG SGSGSGSGSGSGYSGSGSGSGSGSGSGSGYSGSGSGSGSGS |
| 0.1$Y_{20}$ | GSGSYGSGSGSGSGYSGSGS |
| 0.1$Y_{40}$ | GSGSYGSGSGSGSGYSGSGSGSGSYGSGSGSGSGYSGSGS |
| 0.1$Y_{60}$ | GSGSYGSGSGSGSGYSGSGSGSGSYGSGSGSGSGYSGSGSGSGSYGSGSGSGSGYSGSGS |
| 0.1$Y_{120}$ | GSGSYGSGSGSGSGYSGSGSGSGSYGSGSGSGSGYSGSGSGSGSYGSGSGSGSGYSGSGSGSGSYGSGSGSGSGYSGS GSGSGSYGSGSGSGSGYSGSGSGSGSYGSGSGSGSGYSGSGS |
| 0.15$Y_{20}$ | GSGYSGSGSYGSGSGSYGSG |
| 0.15$Y_{40}$ | GSGYSGSGSYGSGSGSYGSGSGSYGSGSGSYGSGSGYSGS |
| 0.15$Y_{60}$ | GSGYSGSGSYGSGSGSYGSGSGSYGSGSGSYGSGSGYSGSGSGYSGSGSYGSGSGSYGSG |
| 0.15$Y_{120}$ | GSGYSGSGSYGSGSGSYGSGSGSYGSGSGSYGSGSGYSGSGSGYSGSGSYGSGSGSYGSGSGYSGSGSYGSGSGSYGS GSGSYGSGSGSYGSGSGYSGSGSGYSGSGSYGSGSGSYGSGS |
| 0.2$Y_{20}$ | GSYGSGSYGSGSYGSGSYGS |
| 0.2$Y_{40}$ | GSYGSGSYGSGSYGSGSYGSGSYGSGSYGSGSYGSGSYGS |
| 0.2$Y_{60}$ | GSYGSGSYGSGSYGSGSYGSGSYGSGSYGSGSYGSGSYGSGSYGSGSYGSGSYGSGSYGS |
